# Dissecting heterogeneous cell populations across drug and disease conditions with PopAlign

**DOI:** 10.1101/421354

**Authors:** Sisi Chen, Jong H. Park, Tiffany Tsou, Paul Rivaud, Emeric Charles, John Haliburton, Flavia Pichiorri, Matt Thomson

**Affiliations:** Division of Biology and Biological Engineering, California Institute of Technology. Pasadena, California, 91125, USA; Department of Molecular and Cell Biology, University of California - Berkeley, Berkeley. CA 94720, USA; Augmenta Bioworks Inc. 3475 Edison Way, Suite K, Menlo Park. CA 94025; Beckman Center for Single-cell Profiling and Engineering. Pasadena, California, 91125, USA; Department of Hematologic Malignancies Translational Science, City of Hope. Monrovia, California, 91016, USA

## Abstract

Single-cell measurement techniques can now probe gene expression in heterogeneous cell populations from the human body across a range of environmental and physiological conditions. However, new mathematical and computational methods are required to represent and analyze gene expression changes that occur in complex mixtures of single cells as they respond to signals, drugs, or disease states. Here, we introduce a mathematical modeling platform, PopAlign, that automatically identifies subpopulations of cells within a heterogeneous mixture, and tracks gene expression and cell abundance changes across subpopulations by constructing and comparing probabilistic models. Probabilistic models provide a low-error, compressed representation of single cell data that enables efficient large-scale computations. We apply PopAlign to analyze the impact of 40 different immunomodulatory compounds on a heterogeneous population of donor-derived human immune cells as well as patient-specific disease signatures in multiple myeloma. PopAlign scales to comparisons involving tens to hundreds of samples, enabling large-scale studies of natural and engineered cell populations as they respond to drugs, signals or physiological change.

## Introduction

All physiological processes in the body are driven by heterogeneous populations of single cells [1, 2, 3]. Single-cell measurement technologies can now profile gene expression in thousands of cells from heterogeneous cell populations across different tissues, physiological conditions, and disease states. However, converting single cell data into models that provide a population-level understanding of processes like an immune response to infection or cancer progression remains a fundamental challenge. All human tissues contain many different subpopulations of cells, and each subpopulation can undergo distinct changes in gene expression and cellular abundance in response to signals, drugs, or environmental conditions. New conceptual and mathematical frameworks are required to model and track the changes that occur within distinct subpopulations of cells within a heterogeneous tissue as they respond to perturbations or succumb to disease.

In this paper, we introduce a computational framework, PopAlign, that identifies, aligns, and tracks subpopulations of single cells within a heterogeneous cell population profiled by single cell mRNA-seq [2, 4, 5, 6]. Mathematically, PopAlign constructs a probabilistic model of each cell population across a series of samples. PopAlign (a) automatically identifies and models subpopulations of cells (b) aligns cellular subpopulations across experimental conditions (signaling, disease) and (c) quantifies changes in cell abundance and gene expression for all aligned subpopulations of cells.

The key conceptual advance underlying PopAlign is representational: we model the distribution of gene expression states within a heterogeneous cell population using a probabilistic mixture model that we infer from single cell data. PopAlign identifies and represents subpopulations of cells as independent Gaussian densities within a reduced gene expression space. PopAlign, then, makes quantitative statistical alignments between subpopulations across samples, and thus enables targeted and quantitative comparisons in gene expression state and cellular abundance. Probabilistic modeling is enabled by a novel low dimensional representation of cell-state in terms of a set of gene expression features learned from data [7, 8, 9]. Unlike PopAlign, geometric methods based on global cell clustering [10, 11] do not provide a natural language for mathematically representing a subpopulation of cells or statistical metrics for quantifying shifts in population structure across experimental samples.

Critically, PopAlign fulfills a fundamental need for comparative analysis methods that can scale to hundreds of experimental samples. Fundamentally, PopAlign runtime scales linearly with the number of samples because computations are performed on probabilistic models rather than on raw single cell data. Probabilistic models provide a reduced representation of single cell data, reducing the memory footprint of a typical 10,000-cell experimental sample by 50 − 100x. Further, downstream computations including population alignment are performed on the models themselves, often reducing the number of computations by an order of magnitude. By contrast methods based on extraction of geometric features (clusters) from single cell data either by clustering (Louvain) or tSNE rely on pairwise computations between individual cells, which is compute-intensive, and requires storing of many raw single cell data sets in memory.

We assess the accuracy and generality of PopAlign using twelve datasets from a mouse tissue survey (Tabula Muris) [12] as well as new experiments on human peripheral blood cells, including a screen of immunomodulatory drugs and a comparison of healthy patients to disease (multiple myeloma). We show that PopAlign can identify and track cell-states across a diverse range of tissues, drug perturbation experiments, and human disease states. The probabilistic models have high representational accuracy and identify biologically meaningful cell-states from data. We performed an experimental screen of 40 immunomodulatory compounds applied to primary human immune cells, and used PopAlign to discover the biggest hits at a population-level and also for specific cell types within the mixture. Finally, we used PopAlign to extract general and treatment-specific signatures of disease progression from multiple myeloma patient samples. Moving forward, PopAlign sets the stage for the analysis of large-scale experimental screens of drugs and genetic perturbations on heterogeneous cell populations extracted from primary human tissue samples.

## Key Contribution

- Probabilistic modeling of cell populations is key conceptual and practical advance that enables multi-scale analysis of single cell datasets across samples, subpopulations, and individual cells.
- Application of method to data sets from mouse tissues and primary human cells demonstrates accuracy of models and ability to track cell-state specific gene expression changes in response to drugs and disease states.

## Results

### PopAlign represents heterogeneous cell populations with probabilistic mixture models

We develop a mathematical and computational framework (PopAlign) that (i) identifies and aligns cell-states across paired populations of single cells (a reference population and a test population), and then (ii) quantifies shifts in cell-state abundance and gene expression between aligned populations (Fig. 1). The method has three steps: probabilistic mixture model construction, model alignment, and parameter analysis. PopAlign can be applied to analyze gene expression and population structure changes in heterogeneous populations of cells as they respond to signals, drugs, and disease conditions.

**Figure 1.**
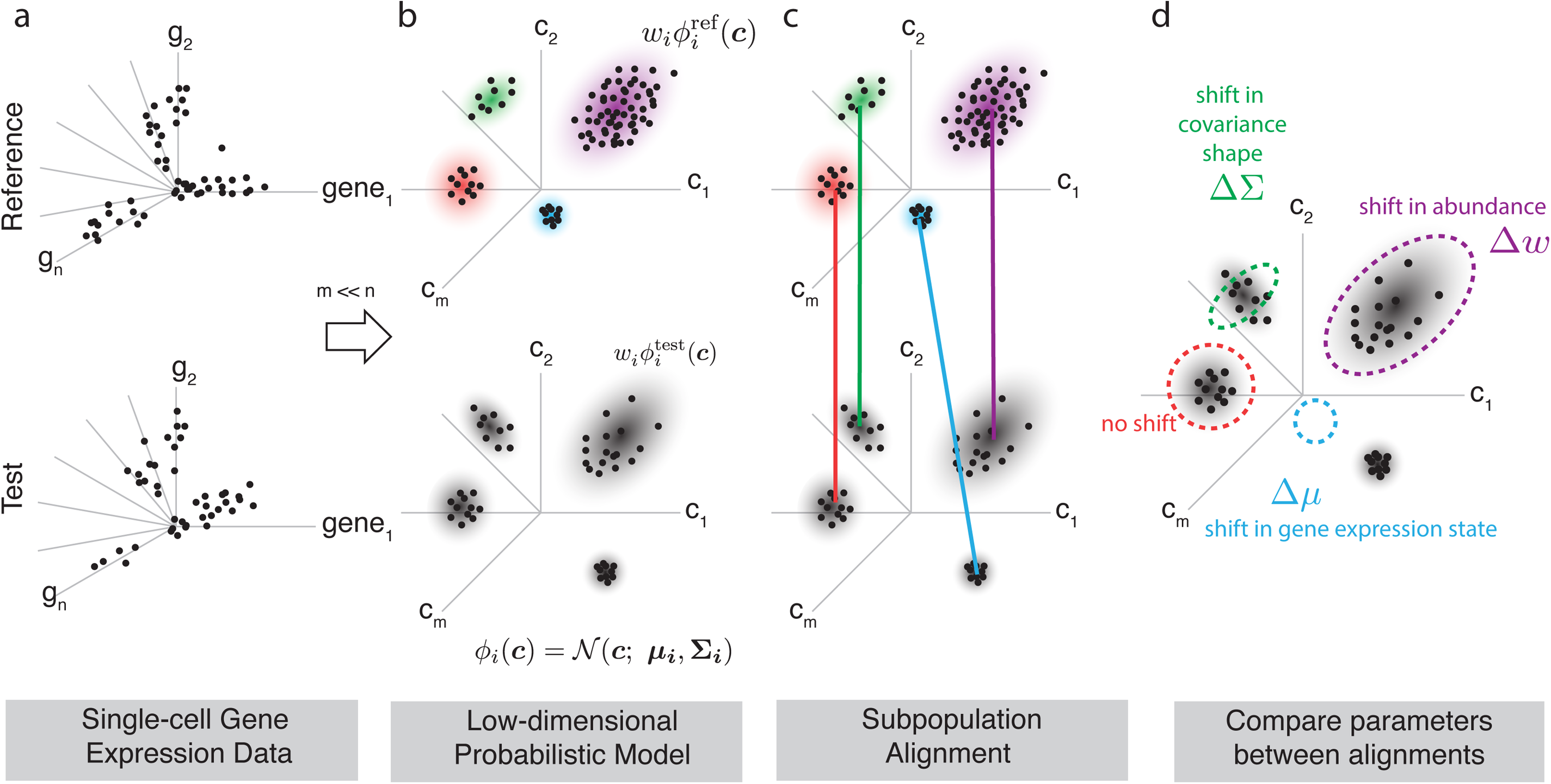
Summary of PopAlign framework. PopAlign provides a scalable method for deconstructing quantitative changes in population structure including cell-state abundance and gene expression across many single cell experimental samples. (a) Users input PopAlign single-cell gene expression data from a ‘Reference’ sample, and at least one ‘Test’ sample, which are each a collection of n-dimensional gene expression vectors ***g***, shown as single dots. (b) For each sample, PopAlign estimates a low-dimensional probabilistic model that represents the distribution of gene expression states as a mixture of local Gaussian densities *ϕ*_*i*_ with parameters encoding subpopulation abundance (*w*_*i*_), mean gene expression state (*µ*_*i*_), and population spread (Σ_*i*_). PopAlign reduces the dimensionality of the input data by representing each gene expression vector as set of *m* gene expression features (*m* = 10 − 20), thus representing each cell as an *m*-dimensional vector of coefficients ***c***. (c) Each 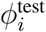 in the test population is aligned to the closest 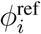 in the Reference sample by minimizing Jeffrey’s divergence. (d) Following alignment, the parameters of aligned subpopulation pairs are compared to identify subpopulation-specific shifts in cellular abundance Δ*w*, shifts in mean gene expression state Δ*µ* and shifts in subpopulation shape ΔΣ.

We consider two populations of cells, a test and a reference population, (***D***^test^ and ***D***^ref^), that are profiled with single cell mRNA-seq (Fig. 1a). Profiling of each population generates a set of gene expression vectors, e.g. 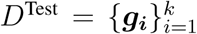 where ***g*** = (*g*_1_, *g*_2_, …, *g*_*n*_), is an *n* dimensional gene expression vector that quantifies the abundance of each mRNA species in single cell ***g*** and *k* is the number of profiled single cells.

To compare the reference and test cell populations, we first, construct a probabilistic model of the gene expression distribution for each set of cells (Fig. 1b). The high dimensional nature of gene expression (*n* ∼ 20, 000) space makes the inference and interpretation of probabilistic models challenging. Therefore, we represent each cell, not as a vector of genes, but as a vector of gene expression programs or gene expression features that are extracted from the data, so that each single cell is represented as a vector ***c*** = (*c*_1_, *c*_2_ … *c*_*m*_) of *m* feature coefficients, *c*_*i*_, which weight the magnitude of gene expression programs in a given cell(See Methods - Extraction of gene feature vectors using matrix factorization). We extract these gene features using a particular matrix factorization method called orthogonal non-negative matrix factorization (oNMF) that produces a useful set of features because all vectors are positive and composed of largely non-overlapping genes (See SI Fig. 1b and 1g). This allows us to naturally think of a cell’s transcriptional state as a linear sum of different positive gene expression programs.

Following dimensionality reduction, for a given cell population, we think of cell states as being sampled from an underlying joint probability distribution over this feature space, *P* (***c***), that specifies the probability of observing a specific combination of gene expression features/programs, ***c***, in the cell population. We estimate a probabilistic model, *P* ^test^(***c***) and *P* ^ref^(***c***), for the reference and test cell populations that intrinsically factors each population into a set of distinct subpopulations each represented by a Gaussian probability density (density depicted as individual ‘clouds’ in Fig. 1b):

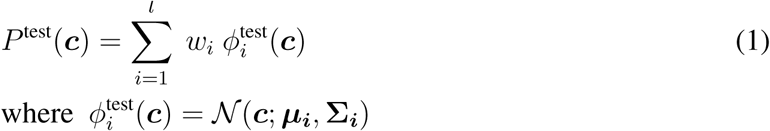

where 𝒩(***c***; *µ*_*i*_, Σ_*i*_) are multivariate normal distributions with weight *w*_*i*_; centroids ***µ***_***i***_ and covariance matrices **Σ**_***i***_. The distributions 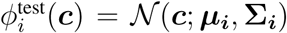, mixture components, represent individual subpopulations of cells; *l* is the number of Gaussian densities in the model. We estimate the parameters of the mixture model (***µ***_***i***_, **Σ**_***i***_, *w*_*i*_) from single cell data using the expectation-maximization algorithm [13, 6] with an additional step to merge redundant mixture components to compensate for fitting instabilities (See Methods - Merging of redundant mixture components).

The parameters associated with each Gaussian density, (***µ***_***i***_, **Σ**_***i***_, *w*_*i*_), have a natural correspondence to the biological structure and semantics of a cellular subpopulation. The relative abundance of each subpopulation corresponds to the weight *w*_*i*_ ∈ [0, 1]; the average cell gene expression state of each subpopulation corresponds to the (*m* dimensional) Gaussian centroid vector ***µ***_***i***_, and the shape or spread of the subpopulation is captured by the covariance matrix **Σ**_***i***_. Intuitively, the local Gaussian densities provide a natural ‘language’ for comparisons between samples. Each Gaussian is a region of high density in gene feature space, and we compare cell populations by asking how the density of cells shifts across experimental conditions.

### Statistical alignment of cellular subpopulations between samples

To compare the test and reference models, we ‘align’ each mixture component in the test population model, 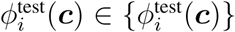, to a mixture component, 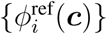, in the reference population model (Fig. 1c). Alignment is performed by finding the ‘closest’ reference mixture component in gene feature space. Mathematically, to define closeness, we use Jeffrey’s divergence, a statistical metric of similarity on probability distributions. We chose Jeffrey’s divergence over other metrics because it is symmetric while also having a convenient parametric form (see Methods)

Specifically, for each 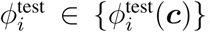, we find an 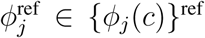, the closest mixture in the reference set:

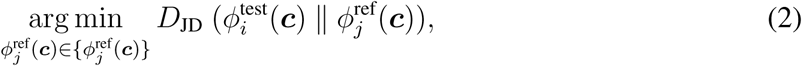

where the minimization is performed over each 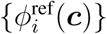 in the set of reference mixtures, and *D*_*JD*_ is the Jeffrey’s divergence (14). Intuitively, for each test mixture, we find the reference mixture *ϕ*_*j*_ that is closest in terms of position and shape in feature space. For each alignment, we can calculate an explicit p-value from an empirical null distribution *P* (*D*_*JD*_) that estimates the probability of observing a given value of *D*_*JD*_ in an empirical set of all subpopulation pairs within a single cell tissue database (See Methods - Scoring alignments).

### Tracking cell-state shifts through mixture model parameters

Following mixture alignment, we analyze quantitative differences in mixture parameters between the reference and test models to track shifts in gene expression state, gene expression covariance, and cellular abundances across the identified subpopulations of cells(Fig. 1d). Mathematically, for each aligned mixture pair, 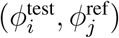 with parameters 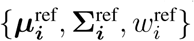 and 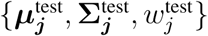, we calculate:

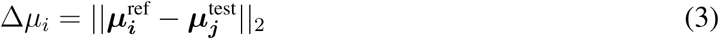

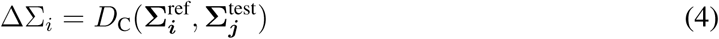

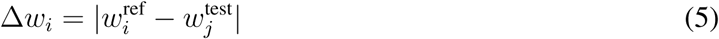

where Δ*µ*_*i*_ measures shifts in mean gene expression; Δ*w*_*i*_ quantifies shifts in cell-state abundance; ΔΣ_*i*_ quantifies shifts in the shape of each mixture including rotations and changes in gene expression variance (see Methods) [14]. We calculate these shifts in parameters for all mixture pairs to assess the impact of drug perturbations or environmental changes on the underlying cell population.

### PopAlign identifies and aligns cell-states across disparate mouse tissues

To test the accuracy and generality of PopAlign, we first constructed and aligned probabilistic models across a wide range of mouse tissues from a recent public study (Tabula Muris) [12]. The Tabula Muris study contains single cell data collected from 12 different tissue samples with ∼ 40, 000 cells total.

For all tissues analyzed, the probabilistic mixture models produce an accurate and interpretable decomposition of the underlying cell states (SI Fig. 3). Accuracy of the models can be assessed by comparing the synthetic (model generated) data to raw experimental data held out from model training. PopAlign models generate synthetic data that replicates the geometric structures and statistical variations found in the tissue data in tSNE or PCA plots with quantitative error of ∼12% (See methods; Fig. 2; 3a,b; SI Tissues; see methods).

**Figure 2.**
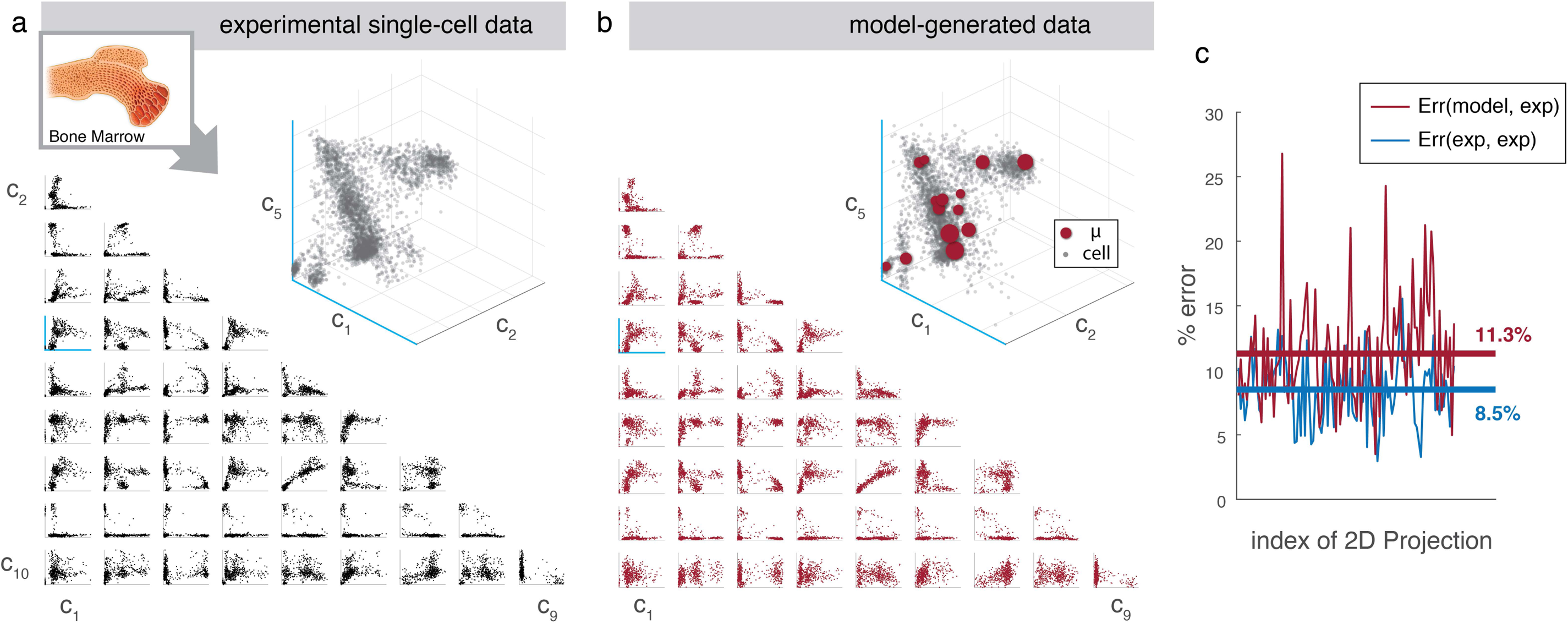
PopAlign models represent experimental data with high qualitative and quantitative accuracy. (a) Experimental data for ∼ 3600 bone marrow cells projected into an *m* = 15 dimensional gene feature space. 2D plots show single cells projected along gene feature pairs (*c*_*i*_, *c*_*j*_), and a single selected 3D projection (inset) is shown. Blue axis denote shared axis between 2D and 3D plot. (b) Model generated data for the same 2D and 3D feature space projections shown in (a). In the 3D projection (inset), each maroon circle denotes the centroid (*µ*) of a Gaussian mixture component where the circle radius is proportional to *w*_*i*_, the mixture weight. In all cases, the model generated data replicates the qualitative geometric structures in the experimental data. (c) Model error across 2D projections from (a) and (b) quantified by analyzing percent deviation in point density for experimental vs model generated data. Quantification of error is performed using a numerical error metric based on binning the 2D projections (See Methods - Analysis of model error). The quantitative error in model-generated is on average 11.3% (red line) compared with 8.5% error when comparing random sub-samples of experimental data (blue line).

**Figure 3.**
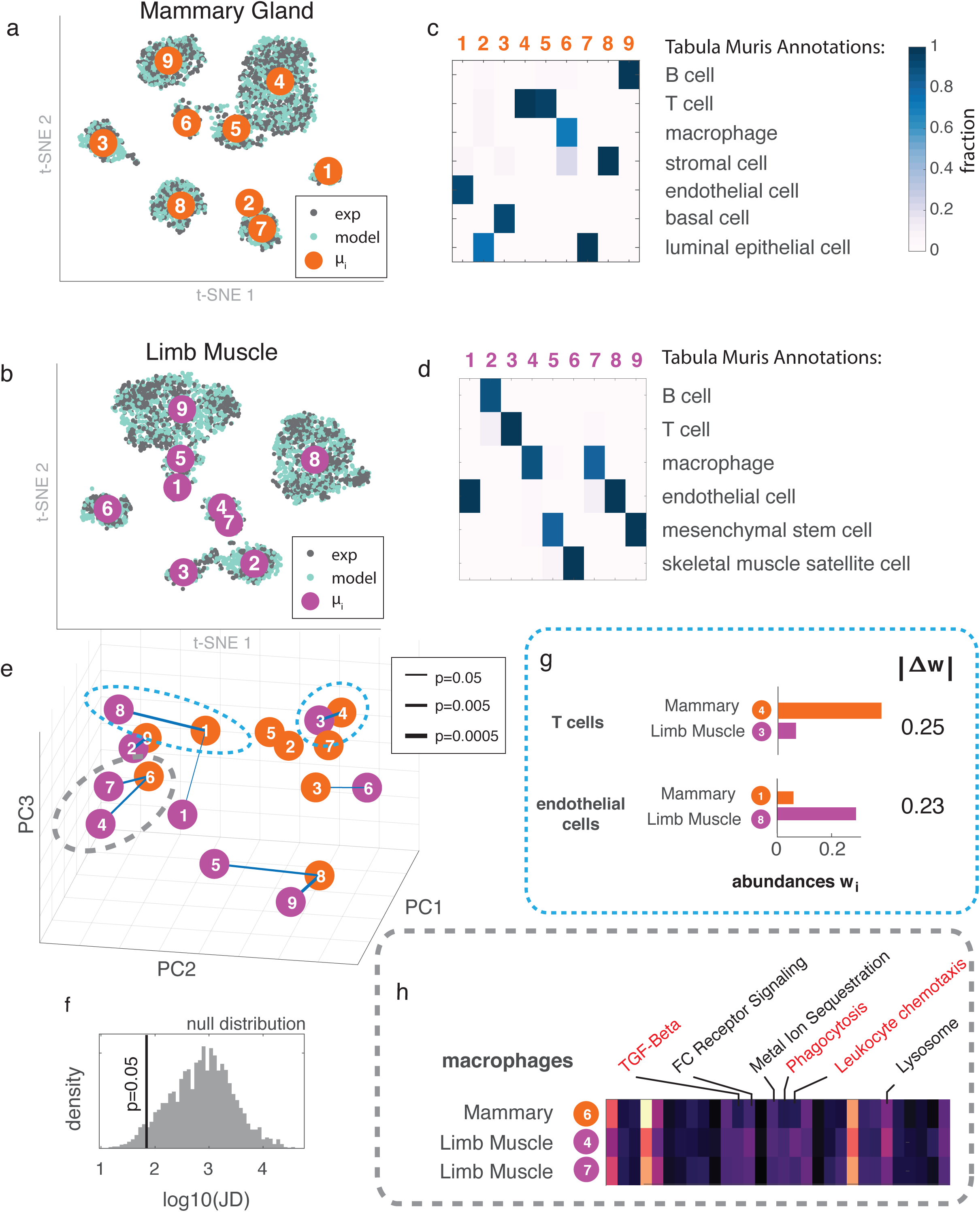
Probabilistic models identify, align, and dissect cellular subpopulations across disparate tissues. Experimental single-cell data (black) for two tissues, mammary gland (a) and limb muscle (b), are plotted together with PopAlign model-generated data (teal) using a 2D t-SNE transformation. Both experimental datasets contain ∼ 3600 cells. For each tissue, mixture model centroids (*µ*) are indicated as numbered disks. (c-d) For models from both tissues, mixture components (x-axis) are scored using cell type annotations supplied by Tabula Muris (y-axis). Each cell of the heatmap represents the percentage of cells associated with each mixture component that have a specific cell type label. Columns (but not rows) sum to 1. (e) Alignments between mixture component centroids (*µ*) from the reference population (Mammary Gland) and the test population (Limb Muscle) are shown as connecting lines. All m-dimensional *mu* vectors are transformed using principal components analysis (PCA) and plotted using the first 3 PCS. Width of each line is inversely proportional to the p-value associated with the alignment (see Legend). (f) Null distribution of Jeffrey’s divergence used to calculate p-values. Jeffrey’s divergence was calculated for all possible pairs of mixture components from models of all tissues from Tabula Muris. (g) We rank aligned subpopulations in terms of maximum Δ*w* and show top two pairs. These subpopulations are identified as T cells and endothelial cells, and are highlighted using a blue dotted line in (e). We find that T cells are highly abundant in Mammary gland while endothelial cells are highly abundant in muscle. (h) Comparing subpopulation centroids (*µ*) for macrophages in terms of gene expression features. Macrophages in Mammary gland and Limb Muscle share common features (black font), but also have tissue-specific features (red font). Corresponding alignments are highlighted in (e) with a gray dotted line.

In addition to providing an accurate representation, the mixture models decompose the cell populations into a biologically interpretable set of cellular subpopulations represented by individual *ϕ*_*i*_(***c***), the mixture components (Fig. 3c,d). The PopAlign mixture components, {*ϕ*_*i*_(***c***)} commonly contain cells of a single cell ‘type’ as defined by labels supplied by the Tabula Muris project. In example tissues, PopAlign extracts known tissue resident cell-types including (Fig. 3c,d) basal cells, luminal cells, macrophages, and T-cells (in mammary gland) and skeletal muscle cells, mesenchymal stem cells, endothelial cells, and macrophages (in limb muscle). Broadly, across all tissue models, 70% of the mixture components classified for a single cell-type provided by Tabula Muris (SI Fig. 3).

Through alignment of model components across tissues, PopAlign enables high-level comparisons of tissue composition. By aligning Mammary Gland to Limb Muscle (Fig. 3e), we identified ‘common’ cell-types between the two tissues including B-cells (p=0.0006), T-cells (p=0.001), endothelial cells (p=0.0013), and macrophages (p=0.004, 0.0076) (SI Fig. 4), and also revealed tissue scale differences in relative abundance. T-cells are highly prevalent (*w* = .3 in the mammary gland but rare in the limb muscle *w* = .05) (Fig. 3g); endothelial cells are highly abundant in the limb muscle (*w* = .32), but rare in the mammary gland (*w* = .06) (Fig. 3g). Between shared cell types, such as macrophages, we reveal common programs such as FC-receptor Signaling and Lysosome, as well as tissue-specific gene expression programs such as TGF-Beta, Phagocytosis, and Leukocyte Chemotaxis(Fig. 3h). PopAlign can, thus, give insight into the underlying composition of a tissue, shedding light onto principles of tissue organization with respect to tissue function.

**Figure 4.**
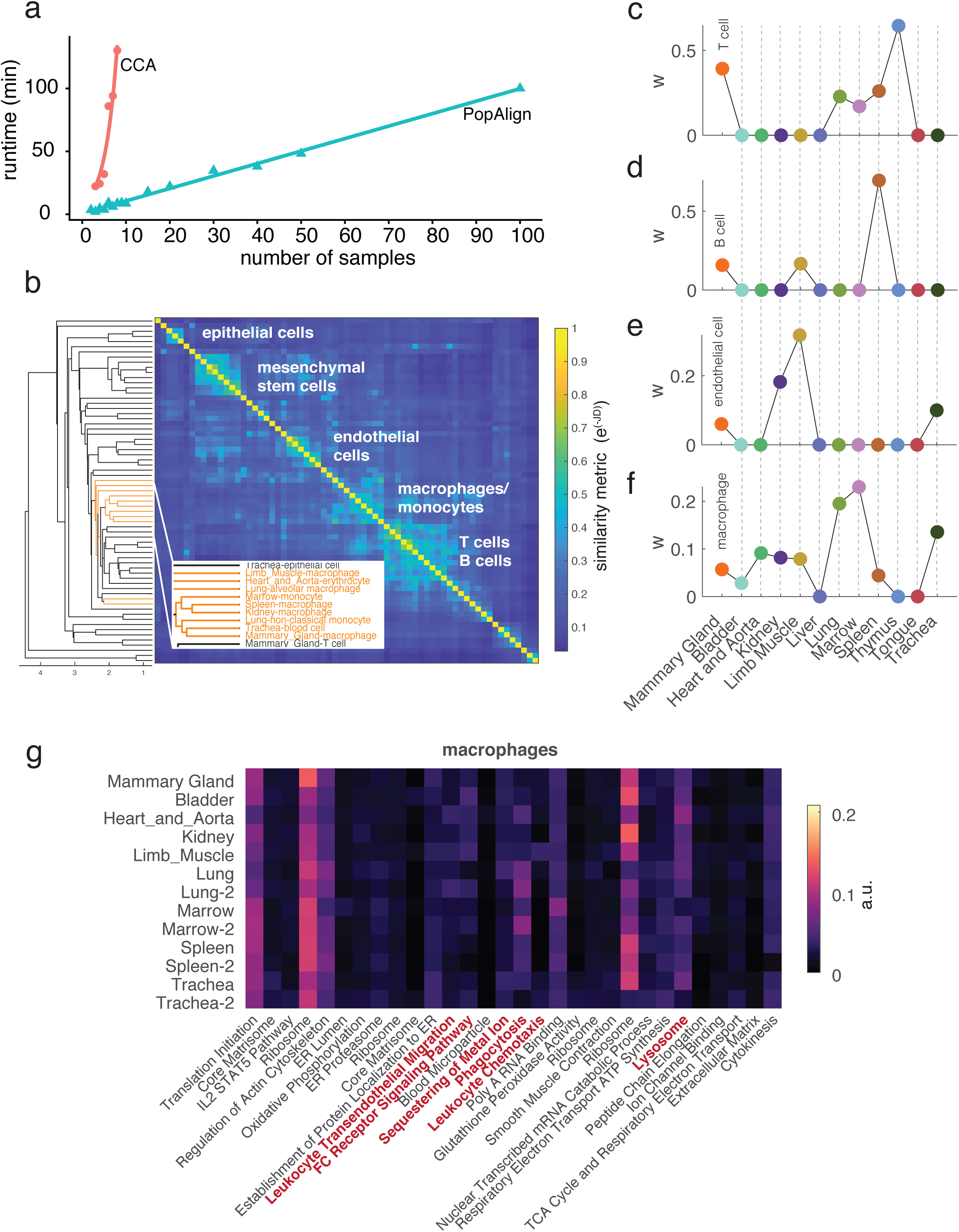
PopAlign can perform global comparisons of cell states across dozens to hundreds of experimental samples. (a) Computational runtime versus number of samples for PopAlign (blue) vs Seurat’s CCA-based alignment method (red). PopAlign scales linearly with the number of samples, while CCA scales exponentially and encounters an out-of-memory error when applied to ¿ 8 samples. Samples are bootstrapped from all 12 samples of the mouse tissue survey Tabula Muris. Benchmarking tests performed on typical workstation (8 cores, 64GB RAM). (b) Heatmap of a pairwise similarity metric between subpopulations from all 12 tissues demonstrates PopAlign can identify cogent cell-type specific clusters even when applied on very disparate tissue types. The similarity metric is defined as exp(-JD) where JD is the Jeffrey’s Divergence between two subpopulations. Inset highlights subpopulations clustered as macrophages, displaying tissue and cell type labels extracted from Tabula Muris annotations. (c-f) Models for all tissues are aligned to a reference model (Mammary Gland) and corresponding abundances (w) are plotted for selected subpopulations classified as (c) T cells (d) B cells (e) endothelial cells (f) macrophages. (g) Mean gene expression state () for macrophage across all tissues show variation in key immune pathways (highlighted in red).

### PopAlign can perform global comparisons of cell state across tens to hundreds of samples

We tested the ability of PopAlign to compare large numbers of samples, using synthetic collections of samples bootstrapped from Tabula Muris data survey. We found that PopAlign runtime scales linearly with sample number and can analyze 100 samples in approximately 100 minutes on a typical workstation with 8 cores and 64GB RAM (Fig. 4a). By first building models, PopAlign front-loads the computation to produce a low-error (Fig. 2) representation of the data that achieves a 50-100x reduction in the memory footprint. Memory efficiency speeds up downstream tasks, such as the calculation of pairwise divergences between subpopulations (Fig. 4b) necessary for aligning them across samples.

Applying PopAlign to compare all 12 tissues of Tabula Muris shows the method is general across many types of experiments, including comparisons of disparate tissues that do not contain overlapping populations. PopAlign achieves generality because it aligns subpopulations by performing a local computation for each test subpopulation (i.e. the minimization of Jeffrey’s Divergence relative to reference subpopulations), that can be accepted or rejected by a hypothesis test. Other methods for comparing samples across experiments essentially perform batch correction to align multiple datasets, before pooling data and jointly identifying clusters [10, 11]. These alignment methods discover a global transformation to bring together cells that are known or inferred to be transcriptionally similar. Not only are these alignment methods computationally slow, scaling exponentially with cell/sample number (Fig. 4a), but they also require overlapping populations or force them to overlap, thus limiting the generality of the approach.

In the 12-sample Tabula Muris comparison, PopAlign uncovered meaningful signatures of cell distributions and gene expression patterns that reflect and expand upon known biology. For example, we found that T cells (Fig. 4c) and B cells (Fig. 4d) are most abundant in organs where they are known to mature developmentally (the thymus [15] and spleen[16] respectively), endothelial cells (Fig. 4e) are most prevalent in highly vascularized tissues (Kidney and Limb Muscle), and macrophages (Fig. 4f) are highly prevalent in the Lung, which accumulates debris and bacteria that must be engulfed and destroyed. The analysis also highlights surprising results, such as the observation that T cells are very abundant in the mammary gland (Fig. 4c). We also found distinct patterns of gene program activation (e.g. Lung macrophages are highly phagocytic) in macrophage populations across tissues (Fig. 4g), consistent with previous reports of functional diversity among macrophages [17]. These results demonstrate that PopAlign is an efficient computational framework for extracting meaningful shifts in abundance and gene expression that scales to large numbers of samples, and is not constrained by requirements for overlapping cell populations between samples.

### PopAlign identifies universal and cell-type specific impacts of drugs

A key application of PopAlign is to study heterogeneous cell populations from the human body as they respond to environmental change, drug treatments, and disease. The human immune system is an important application domain for PopAlign as an extremely heterogeneous physiological system that is central for disease and cell engineering applications [2, 18, 19, 20, 21]. Being able to screen the effects of different drugs on complex immune cell populations, and understand how they affect cell function, is fundamentally important to our ability to design drug therapies for disease treatment. Thus, we performed an analysis of commercially available immunological compounds on human immune cells and used PopAlign to discover how these compounds alter specific cellular subtypes.PopAlign allows us to explore the data hierarchically, first by using quantitative statistical metrics to rank samples and identify interesting drug hits at the population-level, and then by dissecting the impact of these hits on subpopulation composition and gene expression programs.

We performed our screen using 40 drugs (Fig. 5a) from a commercially available compound library (Selleck Chem) on peripheral blood mononuclear cells (PBMCs) from a healthy 22-year old male donor. PBMCs normally contain a mixture of different immune cell types, but our model revealed that blood samples from this particular donor were dominated by monocytes (18%) and T cells (82%) (Fig. 1a).

**Figure 5.**
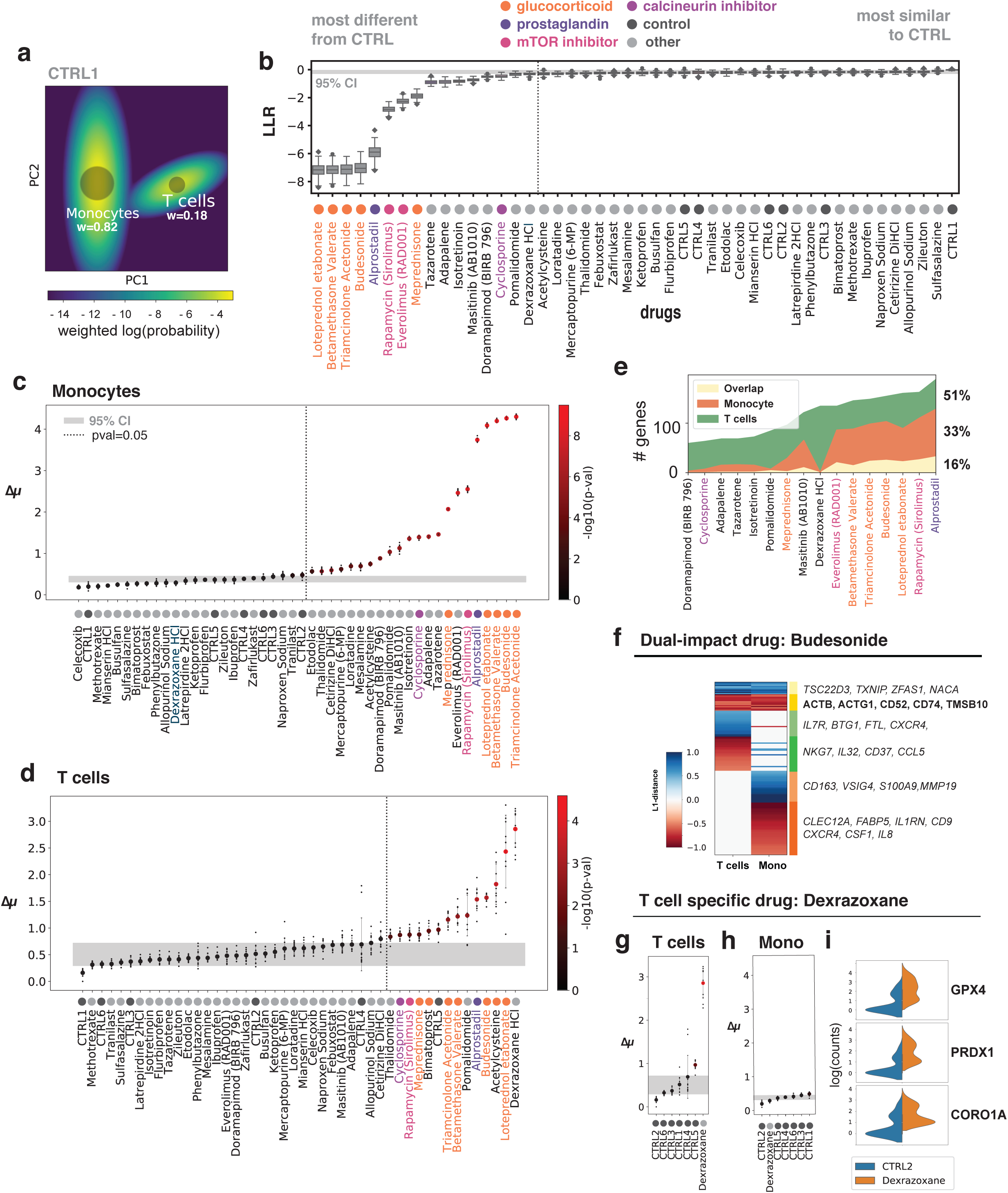
PopAlign identifies universal and cell-type specific impacts of immunomodulatory drugs. (a) Rendering of GMM model for the control sample 1 projected onto the first 2 principal components. Abundance weights (w) are represented by the size of the circle, and supplied as a text label. (b) Ranking of all drugs based on population-level similarity to control population using the log-likelihood ratio metric (LLR). P-values are calculated using an FDR-corrected one-sample t-test of the 6 control replicates against the drug’s mean LLR. Dashed line: p-value = 0.05. (c)-(d) Gene expression shifts (Δ*µ*) for drug-exposed (c) monocyte and (d) T cell subpopulations with respect to their aligned subpopulation in control sample 1. Each small black dot represents a separate bootstrapped model built from a randomly chosen subsample (80%) of the same data. The large dot indicates the mean Δ*µ*, and is colored by - log(p-value). P-values are calculated using an FDR-corrected one-sample t-test testing the 6 control replicates against the drug’s mean Δ*mu*. Gray box: The 95% confidence interval of the control mean. Dashed line: p-value = 0.05. (e) Number of T-cell-specific, monocyte-specific, and overlapping genes across drugs with an LLR p-value *<* 0.05. The maximum percentage of shared genes is 16%. (f) L1-distance metrics are shown for Budesonide, a glucocorticoid that impacts both monocytes and T cells. Differentially expressed genes that overlap between both cell types are denoted in yellow. Up-regulated genes: blue. Down-regulated genes: red. Non-significant L1-distance values: white. (g) T-cell gene expression shifts (Δ*µ*) for dexrazoxane relative to controls. (h) Monocyte gene expression shifts (Δ*µ*) for dexrazoxane relative to controls. (i) Gene expression distributions showing up-regulation of GPX4, CORO1A, and PRDX1 in Dexrazoxane-exposed T cells.

We first identified hits at a high-level by ranking drugs based on how similar the drug-exposed populations are to the unperturbed control populations (6 independent replicates). Statistically, we could define hits as drugs which have a negative log likelihood ratio metric (See Methods - Ranking populations) that lies below the control range (gray box). Within this group, high ranking drugs include a group of glucocorticoids (compounds labeled in orange - Fig. 5b), as well as mTOR inhibitors (pink), alprostadil (a prostaglandin) (purple).

Many immune-regulating drugs are known to be broadly suppressive or activating, but their celltype specific effects are not very well understood. By quantifying and ranking cell type specific shifts, we found that 26 drugs exert significant gene expression shifts (Δ) on monocytes (Fig 5c) while 14 drugs exerted significant effects on T cells (FDR-corrected p-values *<* 0.05) (Fig 5d). Of these drugs, 8 drugs impacted both cell types (Fig 5c,5d, all drugs highlighted in color). Most drugs either did not affect abundances (Δ*w* ≈0) or increased monocyte abundance up to 5% at the expense of T cell abundance (SI Fig 6a,b).

**Figure 6.**
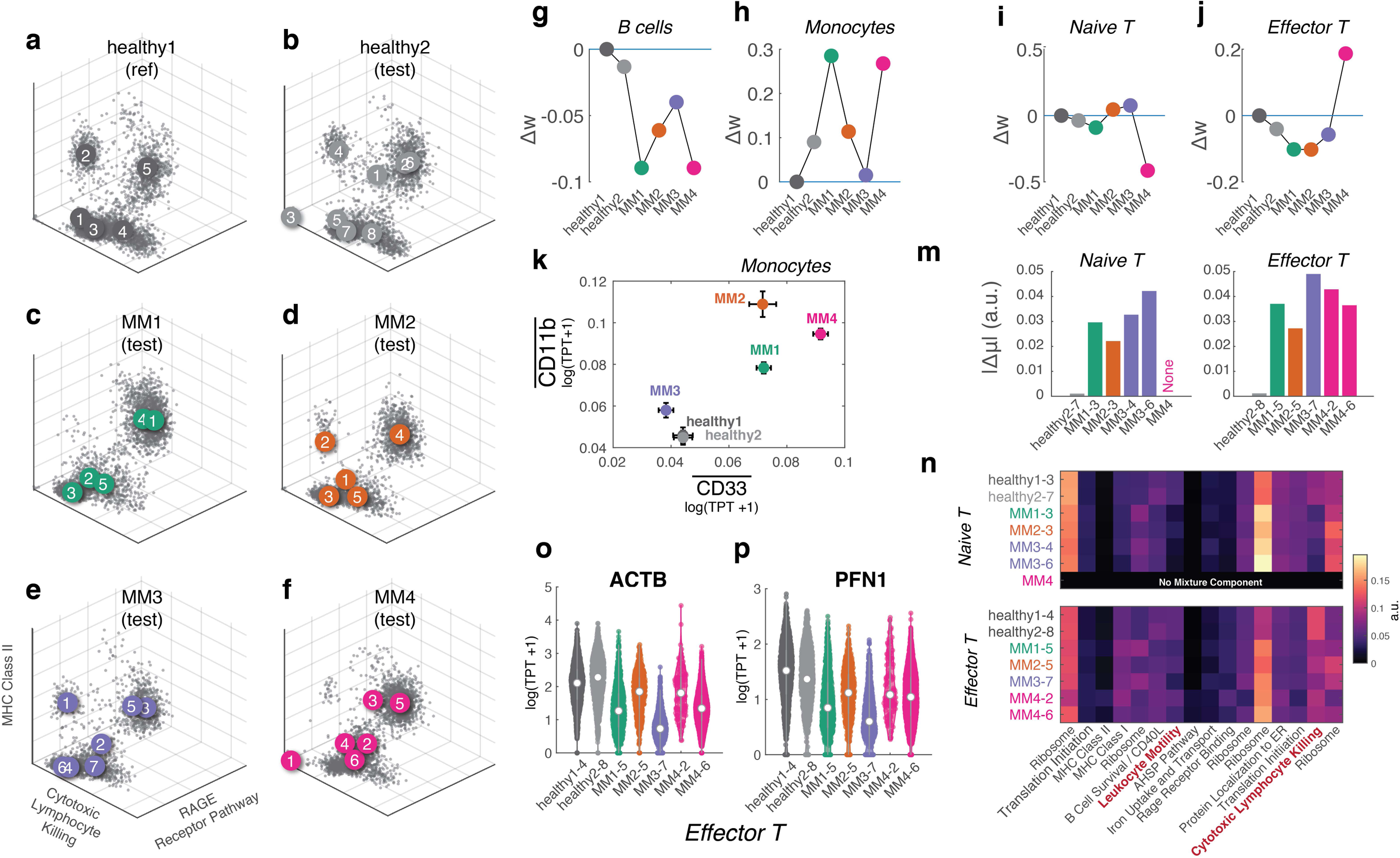
Discovering signatures of disease and treatment in PBMCs from multiple myeloma patients. (a-f) Experimental single cell mRNA-seq data from two healthy donors and four multiple myeloma patients (MM1-4) are projected into 16-dimensional gene feature space. 3D plots show single cells in a subset of three gene features that highlight separation between different immune cell types. Mixture model centroids (*µ*) are indicated as numbered disks. Subpopulations in test samples are aligned to the reference (a) and changes in abundance (Δ*w*) are plotted for (g) B cells, (h) monocytes, i) naive T cells, and j) effector T cells, showing general and patient specific changes. (k) Mean gene expression levels for two markers of myeloid derived suppressor cells (MDSC), CD33 and CD11b, are plotted for all monocyte subpopulations. Error bars denote confidence interval of the mean. (m) |Δ*µ*| for naive T cell and effector T cell populations relative to healthy1. (n) Heatmap of mixture component *µ* vectors in terms of feature coefficients *c*_*i*_ for aligned naive and effector T cells across samples. MM subpopulations exhibit reduced expression of two features (red font): Leukocyte motility and Cytotoxic Lymphocyte Killing. (o) Distribution of beta-actin (ACTB) expression for all effector T cell subpopulations across samples. Violin shows distribution, and mean is denoted by white circle. (p) Distribution of perforin 1 (PFN1) expression for all effector T cell subpopulations across samples. For single gene plots (k), (o), (p), units are in terms of normalized and transformed gene expression (log(g + 1)).

The ability to find the transcriptional impacts of genes that are universal across cell types can reveal important insights into a drugs fundamental mechanisms. In our screen, we discovered that although drug-responsive genes were mostly cell type specific (Fig 5e, Fig), for some drugs, up to 15% of impacted genes were shared between cell types (See Supplementary File 1, which supplies differentially expressed genes for all drugs/cell types). For example, budesonide, up-regulated 11 genes and downregulated 14 genes in both T cells and monocytes (Fig 5f). The overlapping down-regulated genes include many genes associated with actin-based motility - such as actin genes (ACTB, ACTG1), an anti-adhesion peptide (CD52), a myosin interacting protein (CD74) [22] and an actin-sequestering protein (TSMB10) [23]. This result is consistent with earlier observations that glucocorticoids impede T cell polarization and motility [24] and monocyte migratory behavior[25], and suggest that broad leukocyte motility deficits may be partly responsible for the general immunosuppressive effects of glucocorticoids.

Our analyses also allowed us to discover a highly T-cell specific drug, dexrazoxane, which exerted the largest changes on T cell state (mean Δ*µ* = 2.64, p-val = 2.54e-5, Fig. 5g), but no changes in monocytes (mean Δ*µ* = 0.29, p-val = 1, Fig 5h). Dexraxozane did not generate any differentially expressed genes in monocytes (Fig 5e). We found that in T cells, dexrazoxane upregulates many cell survival genes including antioxidant enzymes (GPX4, PRDX1) and CORO1A, which is essential for T cell survival [26] (Fig 5i). Dexrazoxane is normally used as a chemoprotectant agent to reduce toxic side effects of chemotherapy on cardiac tissue [27]. Our finding that dexrazoxane specifically impacts T cells by up-regulating genes that reduce oxidative stress has not been previously reported and could potentially be useful in modulating T cell behavior for other diseases.

PopAlign to allows us to rapidly identify cell-type specific effects of drugs. Identification of the most impactful drugs would be difficult using common visualization approaches like t-SNE (SI Fig. 7) or UMAP, which show qualitative changes (see highlighted conditions), but cannot be readily interpreted because the nonlinear embedding means that changes are not quantifiable. Here, using a small screen of 40 drugs from an immunomodulatory compound library, we were able to use PopAlign to discover universal and cell-type specific mechanisms of drugs, including the observation that glucocorticoids broadly down regulate motility genes and dexrazoxane specifically impacts T cells by upregulating pro-survival genes. Understanding the cell type specific impacts of drugs, which have so far been obscured, will be integral for designing precision therapeutics that have targeted effects within a heterogeneous tissue.

### PopAlign finds general and treatment-specific signatures of multiple myeloma

Given the success of the PopAlign framework in extracting cell-type specific responses in the immune drug response data, we applied the method to study underlying changes in cell state due to a disease process. As a model system, we applied PopAlign to compare human PBMC samples from healthy donors to patients being treated for multiple myeloma (MM). Multiple myeloma is an incurable malignancy of blood plasma cells in the bone marrow. Both the disease and associated treatments result in broad disruptions in cell function across the immune system [28, 29, 30, 31] further contributing to disease progression and treatment relapse. In MM patients, immune cells with disrupted phenotypes can be detected in the peripheral blood[32, 30, 33]. An ability to monitor disease progression and treatment in the peripheral blood could therefore provide a powerful new strategy for making clinical decisions.

We obtained samples of frozen PBMCs from two healthy and four multiple myeloma patients undergoing various stages of treatment (SI Table 1). We profiled *>* 5, 000 cells from each patient, and constructed and aligned probabilistic models to one reference healthy population (Fig. 6a-f).

PopAlign identified several common global signatures in the MM samples at the level of cell-type abundance and gene expression. Across all samples, we find previously known signatures of multiple myeloma including a deficiency in B cells [30, 34, 35], and an expansion of monocyte/myeloid derived cells [32], and critically, new impairments in T-cell functions.

Plotting Δ*w* across all patients, we find high-level changes in subpopulation abundances, which are known to be prognostic of disease progression [33]. We find that all MM patients experience a contraction in B cell numbers (Fig. 4b), and 2 out of 4 see a dramatic expansion (Δ*w >>* .10) of monocytes (Fig. 4c). Changes in T cell levels, however, can be highly variable, with outlier patient MM4 experiencing a large increase in effector T cell (Δ*w* = .2), and a complete elimination of resting T cells (Δ*w* = .2). For this patient, who was receiving a thalidomide-derived drug therapy, these deviations are consistent with thalidomide’s known stimulatory effects on T-cells [36].

Especially in patients with apparently normal abundances (i.e. Δ*w* are small), uncovering subpopulation - specific changes in transcription can point to specific modes of immune dysfunction. We use PopAlign to find that monocyte subpopulations in patients acquire immunosuppressive phenotypes, evidenced by upregulated expression of CD11b and CD33. Both genes are specific markers of myeloid derived suppressor cells[37] which are negative regulators of immune function associated with cancer. By plotting the monocyte-specific mean gene expression values for both CD11b and CD33, we see that all patients except patient MM3 score highly for both MSDC markers. (Fig. 6k). Patients with high MDSC populations typically have a poor prognosis, underscoring the need to monitor MDSC populations in patients.

Importantly, we also find that naive and effector T cells across all multiple myeloma patients have transcriptional defects in pathways essential for T cell function. By plotting Δ*µ*, we show that both populations of T cells experience large mean transcriptional shifts, compared to T cells from our second healthy donor, healthy2 (Fig. 6m). By examining the *µ′s* in terms of gene expression vectors (Fig. 6n), we find that in multiple myeloma, T cells reduce their expression of two key features - Leukocyte Motility, and Cytotoxic Lymphocyte Killing. Surprisingly, the impact on motility is apparent even on the expression of beta-actin (ACTB) (Fig. 6o), a core subunit of the actin cytoskeleton, and which was the top hit in the Leukocyte Motility feature. We find similar declines in the distribution of Perforin 1 (PFN1), a pore-forming cytolytic protein that was found as a top hit in the Cytotoxic Lymphocyte program (Fig. 6p).

Our analysis establishes that we can extract consistent and also patient-specific transcriptional signatures of human disease and treatment response from PBMCs. Interpreting these signatures in the context of disease progression or drug response can provide insight into treatment efficacy and can form the basis of a personalized medicine approach. Our framework enables new applications by providing a highly scalable way of extracting, aligning, and comparing these disease signatures, across many patients at one time.

## Discussion

In this paper, we introduce PopAlign, a computational and mathematical framework for tracking changes in gene expression state and cell abundance in a heterogeneous cell populations across experimental conditions. The central advance in the method is a probabilistic modeling framework that represents a cell population as a mixture of Gaussian probability densities within a low dimensional space of gene expression features. Models are aligned and compared across experimental samples, and by analyzing shifts in model parameters, we can pin-point gene expression and cell abundance changes in individual cell populations.

PopAlign constitutes a conceptual advance over existing single cell analytical methods. PopAlign is explicitly designed to track changes within complex cell populations. Since human diseases like cancer and neurodegeneration arise due to interactions between a wide variety of cell-types within a tissue, population level models will be essential for building a single cell picture of human disease and for understanding how disease interventions like drug treatments impact the wide range of cell-types within a tissue.

Mathematically, existing single cell analysis methods rely on heuristic cluster based analysis to extract subpopulations of cells. Fundamentally, such approaches lack well defined statistical metrics for making comparisons across samples. By conceptualizing a single-cell population as a probability distribution in gene expression space, we define a discrete mathematical object whose parameters can be interpreted, and which can be used to explicitly calculate quantitative statistical metrics for subpopulation alignment. Our probabilistic representation allows us to quickly and scalably learn drug responses even on a complex mixture of cells, in ‘one shot’. This scalability allowed us to analyze data from large-scale drug screen on resting human immune cells, and identify both universal and cell-type specific mechanisms of drugs

In the future, we hope that PopAlign can be used as a part of a work-bench for single cell analysis and treatment of human disease. By applying PopAlign to data sets from the human immune system, we highlight the potential power of PopAlign for identifying drug/signal targets and for deconstructing single cell disease states. PopAlign identified cell-type specific signatures of disease treatment in multiple myeloma patients exposing a potential defect in T-cell activation and motility in three patient samples. This result points to a potential use of PopAlign for guiding treatment interventions by exposing the spectrum of transcriptional states within a diseased tissue and revealing the impact of drug treatments on diseased cell-states as well as the cellular microenvironment and immune cell-types. Such insights could lead to single cell targeting of drug combinations to treat human disease as an essentially population level phenomena.

## Methods

### Mathematical framework

We consider two populations of cells, a reference population, (***D***^test^ and ***D***^ref^), and a test population. Following profiling by single cell mRNA-seq, each population of cells is a set of gene expression vectors, 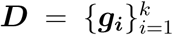 where *k* is the number of cells in the population, and ***g*** = (*g*_1_, *g*_2_, *…, g*_*n*_), is an *n* dimensional vector that quantifies the abundance of each mRNA species. While raw mRNA-seq measurements generate integer valued gene count data, due to measurement noise and data normalization, we consider ***g*** to be embedded in an *n* dimensional Euclidean vector space, gene expression space, ***g*** ∈ ℝ^*n*^. The high dimensional nature of gene expression space poses the key challenge for construction and interpretation of statistical models.

We think of the gene expression vectors, {***g***_***k***_} as being distributed according to an underlying probability density function, *P* (***g***), that quantifies the probability of observing a particular joint gene expression state, ***g*** = (*g*_1_, *g*_2_, *…, g*_*n*_), in a given cell population. Our broad goal is to estimate a statistical model of *P* (***g***),based upon single cell measurements. The model provides a parametric representation of the gene expression density in each condition. Then, we seek to use this representation to track changes in the structure of the cell population across conditions.

In general, the mathematical challenge we face is model estimation in the high dimensional nature of gene expression space. For human cells *n >* 20, 000, and single cell profiling experiments can routinely probe 10,000 cells per sample. The number of parameters in our probabilistic models scales quadratically with *n*, and mixture model learning has data requirements that are exponential in *n* [8]. Therefore, we first reduce the dimensionality of the problem by building models in a common low dimensional space defined by gene expression programs discovered from pooled data across all samples.

#### Data normalization

Single cell gene expression data must be normalized to 1) to account for the variation in the number of transcripts captured per cell and 2) to balance the wide disparity in the scale of values across different genes due to measurement noise and gene drop-out.

The total number of transcripts captured for each single cell can vary from 1000 to 100,000 unique transcripts per cell. Technical variability in reagents and library prep steps can have a large impact on the number of transcripts retrieved per cell. To scale out these differences, we divide each gene expression value *g*_*i*_ by the total number of transcripts and then multiply by a scaling factor *β*.

Additionally, across genes, mean transcript values can span 5 orders of magnitude. Transforming the data using the logarithm brings values across all genes close in scale, while also reducing the skew in the data distributions. The equation for transforming a raw gene expression value *g*_*i*_ (for a single gene) into a normalized gene expression value, 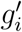 is:

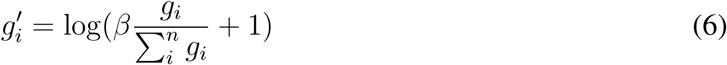

where *n* is the total number of genes, and *β* is a scaling factor, and we add a 1 pseudo-count to each gene expression value. We found that by setting *β* = 1000 to be roughly the median number of total transcript counts in a cell (1000 transcripts), we achieve a smooth transition in the distribution of transformed 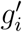 when raw *g*_*i*_ values step from 0 to 1. Gene expression values are thus denoted in units of *log*(*ĝ* + 1) where *ĝ* is cell-normalized and rescaled.

#### Extraction of gene feature vectors with matrix factorization

We circumvent the curse of dimensionality ([7]) by building models in a common low-dimensional space defined by gene expression features or programs. Mathematically, we represent the transcriptional state of each single cell, ***g***, as a linear combination of gene expression feature vectors, {***f***_***i***_}:

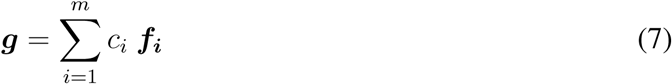

where **f**_**i**_ ∈ ℝ^**n**^ specifies a gene expression feature, and *c*_*i*_ is a coefficient that encodes the weighting of vector *f*_*i*_ in ***g***, the gene expression state of a single cell. The key result in [7] is that a cell’s gene expression state, ***g*** can be represented as a linear combination of *m* gene expression module vectors, ***f***_***i***_ where *m << n*. This insight allows us to construct a low dimensional representation of a cell population and, then, to estimate statistical models within the low dimensional space. [7, 38, 39].

The gene features, {***f***_***i***_} can be extracted using a wide range of matrix factorization and machine learning technique including Singular Value Decomposition, and its matrix factorization relatives like sparse PCA as well as methods like layered neural networks [7]. We use a technique called orthogonal non-negative matrix factorization [40] (oNMF) to define a space of orthogonal gene expression features vectors. Like other linear dimensionality reduction techniques, such as PCA, oNMF factors the original data matrix, ***D***^train^ (SI Fig. 1a) into two matrices ***D*** ≈***F C*** (SI Fig. 1b,c). Factorization occurs through minimization of an objective function with positivity and orthogonality constraints:

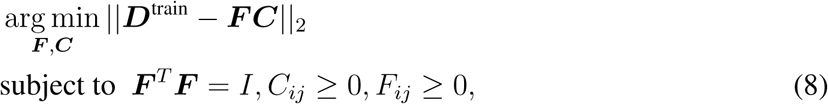

The optimization minimizes the (Frobenius) norm of the difference between the training data, ***D***^train^, and its factored representation ***F C***. The columns of ***F*** contain gene features, ***f***_***i***_. The matrix ***C*** is *m* by *k*, where *k* is the number of single cells in ***D***^train^. Each column of ***C*** encodes the weighting of the *m* gene features across a given single cell. The entries of ***F*** and ***C***; are constrained to be positive, and the columns of ***F*** (the gene features) are constrained to be orthogonal. ***F*** is an *n* by *m* matrix (genes by features). Each column contains *n* weights where each weight corresponds to the weight of a given gene in that feature, ***f***_***i***_.

Standard non-negative matrix factorization has been shown to provide a useful set of features for gene expression analysis because feature vectors have positive entries, and so we can naturally think about the gene expression state of a cell, ***g***, as being assembled as a linear sum of positive gene expression programs.

In PopAlign, we incorporate orthogonality in ***F*** as a secondary constraint to aid interpretation. Empirically, we found that orthogonality aids in interpretation of the features as well as in model construction because the orthogonal gene expression features are interpretable as non-overlapping sets of genes (SI Fig. 1b) that can individually be analyzed by gene set enrichment analysis (SI Fig. 1f)(see Methods - Gene Set Enrichment Analysis). Second, orthogonality tended to force individual cell states to be represented by more than one feature which aided stability during model parameter estimation.

To perform oNMF, we select *m*, the number of features to be extracted through an optimization that balances accuracy and dimensionality explicitly. In oNMF (as opposed to PCA and SVD), *m* is a parameter given to the optimization. Choosing *m* involves balancing the tension between the ‘expressiveness’ in the feature set and its dimensionality. Higher *m* reduces the error in the representation while also breaking up blocks of genes into smaller modules that represent independent gene expression pathways with finer granularity. However, as *m* increases, the typical computational and sampling challenges associated with high dimensionality emerge.

Practically, we balance this tension in PopAlign by constructing a loss function with a penalty that increases with *m*:

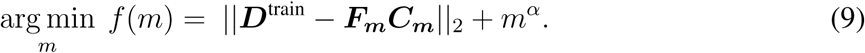

For each value of *m*, we perform oNMF on ***D***^train^ yielding ***F***_***m***_ and ***C***_***m***_, and thus an error ||***D***^train^ − ***F***_***m***_***C***_***m***_ ^2^||. This error is, then, incremented by the term *m*^*α*^ which penalizes higher values of *m* and hence the dimensionality of the feature set. We set *α* = .7 based upon numerical experimentation on model data sets (SI Fig. 2). For any choice of *m*, we can estimate the accuracy of the representation by plotting reconstructed data *FC* (SI Fig. 1d) against normalized data *D* (SI Fig. 1a). The SI shows such plots and the PopAlign software package outputs these plots by default.

Practically, given data sampled a set of cell populations,(eg 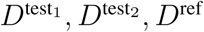) we pool data from all cell populations into a training data set, ***D***^train^, and perform oNMF. If we are analyzing a large number of data sets or sets with many single cells, we generate ***D***^train^ by sampling 500-1,000 cells uniformly at random from the reference and test cell populations and selecting a *m* via ((9)).

Because the feature vectors are not always purely orthogonal, we recast the complete dataset into the feature space using a non-negative least squares. Specifically, for each gene expression profile, ***g***, we find, ***c*** via:

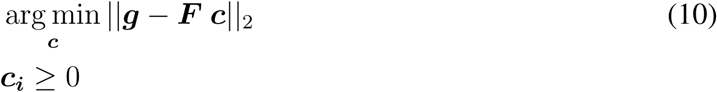

where ***F*** is a fixed feature set learned from oNMF on ***D***^train^. The gene expression vector *g* is thus compressed into a k-dimensional vector *c*_*i*_ that provides a high-level programmatic representation of cell state in terms of gene expression ‘features’. Finally, we interpret the biological meaning of the feature vectors in terms of annotated gene expression programs using gene set enrichment analysis (see Methods - Gene Set Enrichment Analysis). Using matrix factorization, we map a cell population, *D* = {***g***}, from an *n* ∼ 20, 000 dimensional gene expression space into a gene feature space that is often of order 10 − 20 dimensions [7].

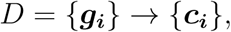

where {***c***_***i***_} are *m ×* 1 dimensional vectors that now represent the cell population in the reduced gene feature space.

#### Gene set enrichment analysis

To interpret the gene features in terms of annotated gene sets, we perform geneset enrichment analysis using the cumulative hypergeometric distribution. We define each feature vector by the collection of genes that have weightings greater than 4 times the standard deviation (*>* 4*σ*). Using this collection of genes, we then calculate the null probability of drawing k genes (*P* (*X > k*)) from a specific annotated gene set using the hypergeometric cumulative probability distribution:

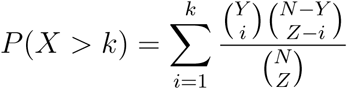

where N is the total number of genes, Z is the number of genes in the feature that are *>* 4*σ*, Y is the number of genes in each annotated gene set, and k is the number of genes that overlap with annotated gene set.

Gene sets are sorted by their associated null probability; the 10 gene sets with the lowest null probabilities are reported for each feature. The gene sets in our dictionary are pulled from GO, KEGG, and REACTOME, and are supplied with our code. SI Fig. 1f shows an example of gene set enrichment results for two features.

#### oNMF error analysis

The error associated with each feature set *F*_*m*_ was assessed by comparing data entries between a cross validation dataset ***D***_***x***_ and its reconstructed matrix ***F***_***m***_***C***_***x***_. We binned the data in ***D***_***x***_ into bins of equal width ∼ 0.1 (in units of log(TPT+1)). We retrieved data values from each bin, (***D***_***x***_)_*i*_, and then plotted their means against the means of corresponding data values in the reconstructed matrix, (***F***_***m***_***C***_***x***_)_*i*_ (SI Fig. 1d). The standard error in each bin is calculated as the mean squared deviation of the reconstructed data from the original data:

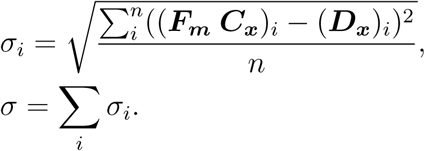

To quantify the amount of dispersion relative to the mean, we also calculate the coefficient of variation for each bin:

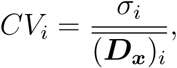

We find empirically that the average CV is ∼ 30 − 35% across all bins for most feature sets.

#### Representing a cell population as a Gaussian Mixture Model in gene feature space

Following the feature based representation, we construct a statistical model of each given cell population within the reduced gene feature space. Mathematically, we have exchanged a probability distribution in gene expression space for a probability distribution in gene feature space:

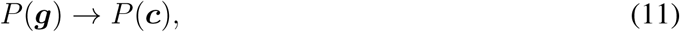

where ***g*** is *n ×* 1 and ***c*** is *m ×* 1, and *m << n*. We can now estimate a statistical model of *P* (***c***).

To account for the heterogeneity of cell-states within a tissue, we model cell-populations using Gaussian mixture models. Gaussian densities provide a natural representation of a transcriptional state in gene expression space which is consistent with measured gene expression distributions as well as empirical models of transcription [41, 42, 43]. Theoretical models of stochastic transcription commonly yield univarite gene expression distributions where mRNA counts are Poisson or Gamma distributed. Normal distributions provide a reasonable approximation to these distributions with a computationally tractable inference procedure.

We represent the cell population as a mixture of Gaussian densities, so that for a given cell population *D*, we construct:

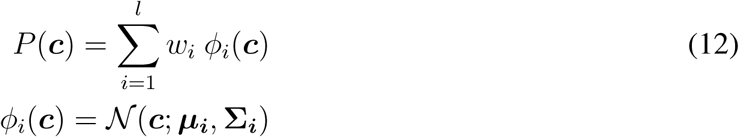

where *P* (***c***) is a mixture of Gaussian densities, 𝒩(***c***; ***µ***_***i***_, **Σ**_***i***_), with centroid, *µ*_*i*_; covariance matrix,, Σ_*i*_; and scalar weighting *w*_*i*_. ***µ***_***i***_ is a vector in the *m* dimensional feature space, and **Σ**_***i***_ is a symmetric *m × m* matrix. *l* is the number of Gaussian mixtures or components in the statistical model. The Gaussian mixture model (GMM) represents a cell populations as a mixture of individual Gaussian densities. Biologically, we think of each density as parameterizing a subpopulation of cells.

We can estimate the parameters ***µ***_***i***_, **Σ**_***i***_, and *w*_*i*_ based upon training data, using maximum likelihood estimation with likelihood function:

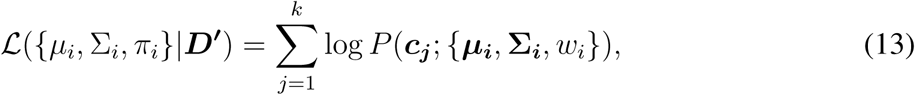

where ***c***_***j***_ are single cell profiles drawn from a cell population ***D***^**′**^ and cast into feature space; *k* is the number of single cells in the cell population ***D***^**′**^. For a given experimental data set, defines a function over the space of model parameters. To select model parameters given data, we can attempt to maximize the value of ℒ. In general for Gaussian Mixture models, likelihood maximum is complicated by the geometry of ℒ which is not concave and can have multiple local and global maxima [44]. ℒ can be maximized approximately using expectation maximization.

Expectation-maximization is a heuristic algorithm that finds (local) maximum likelihood parameters. Although it is known to have fundamental problems - including weak performance guarantees and a propensity to overfit data, new methods [8] place constraints that are invalid for our application (such as shared covariance matrices). We find empirically that the EM algorithm performs well, learning low-error representations of ***c*** (SI Fig. S2), and that we can overcome fitting instabilities by algorithmically merging components. Practically, we perform expectation maximization using sci-kit learn. We regularize the variance of individual mixtures to constrain variance to be non-zero to avoid fitting instabilities. We determine mixtures number through the Bayesian Information Criterion (BIC) which optimizes a trade-off between model complexity and accuracy on training data.

#### Merging of redundant mixture components

One drawback of the EM algorithm is its propensity to fit ‘redundant’ mixture components to the same local density with significant overlap. For our application, this redundancy complicates interpretation and comparisons across samples. We overcome this problem by taking advantage of mixture model properties to algorithmically merge redundant mixtures using the Jeffrey’s divergence.

For multivariate Gaussian distributions, the Jeffrey’s divergence has a closed analytic form.

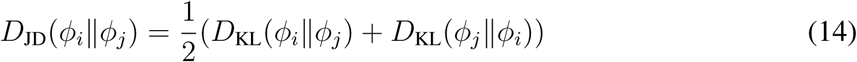

where *ϕ*_0_ and *ϕ*_1_ are two independent components from the same mixture model (12), and *µ* and Σ are their associated parameters.

*D*_KL_ is the Kullback Leibler divergence and has a convenient parametric form for Gaussian distributions:

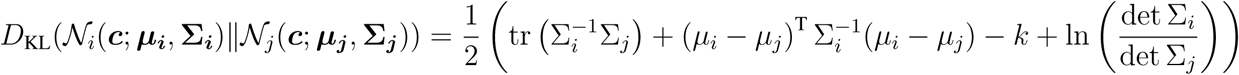

For each mixture model, we iteratively attempt to merge component pairs with the lowest Jeffrey’s divergence and accept mergers that increase the BIC of the model given the data. With each merge step, model parameters for candidate pair are recalculated, and the updated model is accepted or rejected based on the new BIC. Mergers are performed until the first rejection. This procedure removes redundant mixture components from the model.

#### Sampling data from Gaussian mixture models

Given parameters, 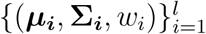 for a given mixture model. We can ‘generate’ synthetic data from the model through a simple sampling procedure. A given model has, *l* mixtures, and we first select a mixture from the set of *l* mixtures with probabilities weighted by *w*_*i*_. Following selection of a mixture, *j*, we use standard methods to draw a ‘point’ in the *m* dimensional feature space from 𝒩_|_(***c, µ***_***j***_, **Σ**_***j***_).

#### Analysis of model error

Model error was assessed by comparing the empirical distributions of model-generated data with experimental data. We performed error analysis within 2D projections of the data, because fully binning a k-dimensional space can be highly memory intensive. Briefly, each 2D projection was binned into 25 bins (5×5). For each bin, the deviation between the model generated data and empirical data was calculated in terms of percent error in each bin. Total error within each projection is calculated as a weighted average of this percent error over all bins in a projection:

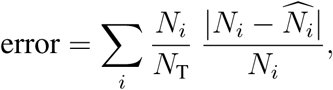

where *N*_*i*_ is the number of experimental data points in bin *i*, and 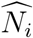 is the number of model generated data points in bin *i. N*_T_ is the total number of experimental data points. This metric weights the per-bin fractional error by the probability density of each binned region in the projection.

### Alignment of models across reference and test populations

To compare the test and reference models, we ‘align’ each mixture component in the test population model, 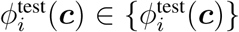, to a mixture component, 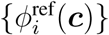, in the reference population model (Fig. 1c). Alignment is performed by finding the ‘closest’ reference mixture in gene feature space. Mathematically, to define closeness, we use, Jeffrey’s divergence, a statistical metric of similarity on probability distributions. Specifically, for each 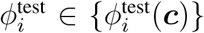, we find an 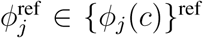, the closest mixture in the reference set:

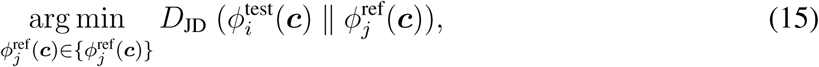

where the minimization is performed over each 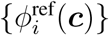 in the set of reference mixtures, and *D*_*JD*_ is the Jeffrey’s divergence (14). Intuitively, for each test mixture, we find the reference mixture *ϕ*_*j*_ that is closest in terms of position and shape in feature space.

Aligning subpopulations using Jeffrey’s divergence incorporates information about the shape and position of the probability distributions (i.e. covariance), while disregarding the relative abundance of each subpopulation.

#### Scoring alignments

For each alignment, we can calculate an explicit p-value from an empirical null distribution *P* (*D*_*JD*_) that estimates the probability of observing a given value of *D*_*JD*_ in an empirical data set of all subpopulation pairs within a single cell tissue database. To assign a p-value to alignments, we calculate the probability of observing two cell-states with a given Jeffrey’s divergence by chance using an empirical null distribution generated from the tissue data set. Specifically, we constructed a mixture model for all tissue pairs in Tabula Muris. Then, we calculate all pair-wise Jeffrey’s divergence scores for all the underlying mixtures. This calculation gives us a global distribution over *D*_JD_ for cell-states in the mouse. This distribution provides a null distribution for typical statistical closeness between cell-states in feature space.

#### Differential gene expression between aligned subpopulations using L1-norm error metric

We can discover up- and down-regulated genes between any two subpopulations by finding genes which have significant shifts in their distributions. We quantify the extent of this shift using a signed L1 distance metric over a discretized domain. For each gene, we calculate two empirical distributions *P*_1_(*a*), *P*_2_(*a*), one for each subpopulation. These distributions are discretized over identical histogram binnings, *a*_*i*_, over the gene’s entire range of expression, *a*_*i*_ ∈ 𝒜. Then we define the signed L1 distance as:

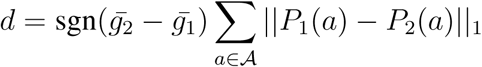

where 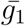 and 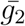 denote the respective means of subpopulations 1 and 2. The sign allows us to distinguish whether a gene is down-regulated in population 2 (negative) or upregulated in population 2 (positive). The signed L1 distance metric ranges from −2 to +2, both of which correspond to completely non-overlapping distributions (which we have not yet seen in gene expression data). In experiments with control samples, we use comparisons between control samples to calculate an empirical cutoff for interpreting an L1-distance as significant. We determine the cutoff as the point at which the fraction of control genes that exceed this value is less than 0.001 (or some user-selected p-value). For the drug screen, this L1 threshold is calculated to be around 0.5. Qualitatively, we see that genes with L1-distances of ¿ 0.5 have distributions which are visually in obvious agreement with the labeled directionality, which holds true across a range of different experimental samples.

### Model interpretation through parameter analysis

Following mixture alignment, we analyze quantitative differences in mixture parameters between the reference and test sample to track shifts in gene expression state, gene expression covariance, and cellular abundances across the identified cell-states in the cell population. Specifically, for each aligned mixture pair, 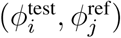 with parameters 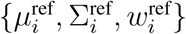 and 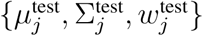 we calculate:

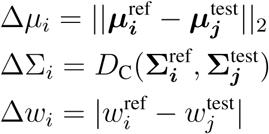

where Δ*µ*_*i*_ measures shifts in mean gene expression; Δ*w*_*i*_ quantifies shifts in cell-state abundance; ΔΣ_*i*_ quantifies shifts in the shape of each mixture including rotations and changes in gene expression variance. For ΔΣ_*i*_, we use the following distance metric on covariance matrices [14]:

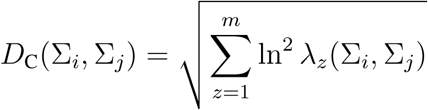

where *λ*_*z*_ is a generalized eigenvalue of **Σ**_***i***_ and **Σ**_***j***_ or a solution to **Σ**_***i***_ *v* = *λ*_*z*_**Σ**_***j***_*v*, and *m* is again the number of gene features.

We calculate these shift parameters for all mixture pairs, and then analyze the shifts to assess the impact of signaling conditions or environmental changes on the underlying cell population.

#### Ranking populations against a control using the log-likelihood ratio metric

The log-likelihood ratio serves as a quantitative measure of how similar perturbed populations are to a control population. To calculate the log-likelihood ratio, we use both the control model (non-perturbed) as well as the perturbed model to compute probability scores for cells sampled from the perturbed sample, and then take the ratio of those probability scores. The LLR is the mean logged ratio between the two probability values, and will be near 0 if the two probabilities are very close (i.e. ratio is near 1) but will be negative if the perturbed sample differs substantially from the control.

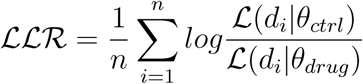

In our drug screen, we observed that the LLR is sensitive to transcriptional changes but this sensitivity is decreased if the affected subpopulation is rare. All of the drugs which have significant LLR (p-value adjusted ¡ 0.05) results in a gene expression shift (delta mu) in either T cells or monocytes. However, drugs that induce cell type-specific changes in very rare cells often do not generate LLRs that are significantly different than the control. For example, Dexrazoxane has an LLR value that barely makes the p-value cut off (Fig 5b), but induces significant gene expression changes in T cells (Fig 5d). There are only 5% T cells in the Dexrazoxane sample (about 40 T cells total), which likely diminishes the T cell impact on the global LLR score.

#### Cell type classification of mixtures using marker genes

For the Tabula muris data set, cells and mixtures were classified using cell-type annotations provided by the study. Mixtures were classified according to the cell-type with the maximum abundance within a given mixture (Fig. 2).

For drug screen experiments, we classified the independent mixtures as T cells (CD3D+), Monocytes (LYZ+), or B cells (CD20+).

For immune cell experiments with healthy and multiple myeloma patients, we classified the independent mixture components as effector T cells (CD3+ / CD57+), naive T-cells(CD3+ / CD28+), erythrocytes (HBB+), canonical monocytes (CD14+ / CD16low), and nonclassical monocytes (CD14low / CD16++) [45] (Fig. 4e). We aligned populations in our test samples - GM-CSF (Fig. 4b) and IFNG (Fig. 3c) - to the reference populations (control) by finding pairs of components that minimize the Jeffrey’s divergence (Fig. 4d).

### Software implementation

PopAlign has been implemented as a Python3 software package. The package requires common scientific computing libraries (numpy, matplotlib, pandas, seaborn, tables, MulticoreTSNE, adjustText) that can be easily installed with pip and our requirements file. A guide on how to get set up and install dependencies is provided on the packages Github page. The software runs on local machines, as well as on Amazon Web Services headless EC2 instances, providing a powerful setting for large-scale analyses. The architecture of this package is built around classes that store experimental samples as objects and provide a set of specific methods to perform tasks such as normalization, dimensionality reduction, model construction, model alignment, and parameter comparison.

### Experimental methods

#### Single-cell RNA-sequencing

Cryopreserved PBMCs (Hemacare) from healthy and multiple myeloma patients were thawed in a 37C waterbath for 2 minutes after which the cells were transferred to a 15mL conical tube. Prewarmed RPMI 1640 was then added to the 15mL conical to a final volume of 10mL and centrifuged for 2 minutes at 300RCF to pellet the cells. Supernatant was removed and cells were resuspended to 1 million cells/mL in RPMI1640 supplemented with 10% FBS and 17,400 cells were loaded into each TENX lane.

#### Sample multiplexing using Multi-Seq

Cryopreserved PBMCs sourced from Hemacare (∼ 50 million cells) were thawed in a 37C waterbath for 2 minutes after which the cells were transferred to a 15mL conical tube. Prewarmed RPMI1640 was then added to the 15mL conical to a final volume of 10mL and centrifuged for 2 minutes at 300RCF to pellet the cells. Supernatant was removed and cells were resuspended in 10mL of RPMI1640 supplemented with 10% FBS and 1% pen/strep. The cell suspension was then plated onto a 100mm low attachment plate and rested in a CO2 incubator at 37C for 16 hours.

After resting, 200,000 cells were loaded into each well of a 96-well plate and exposed to 1 uM of drug in RPMI1640 plus serum. Drugs used were drawn from the Immunology and Inflammation-related library sold by SelleckChem. After 24 hours of exposure, cells were enzymatically dissociated into a single-cell suspension using TrypLE and multiplexed using Multi-seq lipid-modified oligos. [46].

## Supporting information

PopAlign_SI

SI_File1_DrugScreen_DEgenes

## Acknowledgements

Eric Chow, Chris McGinnis, David Patterson, Allan Pool Hermann, Jase Gehring, Members of the Thomson Lab, Beckman Institute Single-cell Profiling and Engineering Center (SPEC).

