## Supplementary material for "Dissecting heterogeneous cell populations across drug and disease conditions with PopAlign": PopAlign_SI

### SI Figure Captions

**SI Figure 1. Orthogonal NMF produces low-error reconstruction** Orthogonal nonnegative matrix factorization factors each a) gene expression data matrix into b) F, a set of orthogonal linear feature vectors, and c) C, a matrix of coefficients. For (b) we order the genes according to their weight in each feature, keeping only genes with greater $> 4\sigma$ for each feature. This ordering highlights the orthogonality of features. (d) The reconstructed data $FC$ is a denoised version of the original data matrix. (e) Data values from the original data matrix are partitioned into bins, and their means are plotted against corresponding data values from the reconstructed data matrix ($FC$). Bin width $\sim 0.1$. Shaded area indicates the standard deviation between original values and reconstructed values within each bin. (See Methods - oNMF error analysis). (f) Histogram of the entries in normalized data matrix D, with log(counts) on y-axis. 99.996% of data values are less than 5, which is also the linear regime of (e). (g) Zoomed in view of white inset from (c), showing specific genes that have high weights for two features, as well as their top gene set results from gene set enrichment analysis.

**SI Figure 2. Choosing dimensionality of feature set** Plot of loss function for different feature sets, built by sweeping over many values of m (x-axis). Feature set with lowest loss (red asterisk) is chosen. This example uses data from Fig. 4. Loss function is a function of the residual error and a penalty on m. For loss function equation, see Methods - Extraction of gene feature vectors with matrix factorization.

**SI Figure 3. PopAlign models for all 12 tissues of Tabula Muris** For each tissue, we plot the following: (upper left) joint t-SNE plot of experimental single-cell data (black), PopAlign model-generated data (teal), and mixture model centroids ($\mu$) as numbered disks, (upper right) Heatmap of the proportion of cells classified by each mixture model component against their annotated labels. Each column sums to 1. (lower) Heatmap of mixture component centroids

($mu$) in terms of their expression level of features. For all tissues, we used a universal 30D feature set determined using sampled data from all tissues within the collection.

**SI Figure 4. Mixture components across Tabula Muris tissues are highly specific for single cell type** Histogram of each mixture component's highest score for an annotated label. These highest scores as taken as the max of each column in Fig 3c,d and analogous cell annotation heatmaps for all tissues in SI Figure 3. Most mixture components score for a single annotated label uniquely.

**SI Figure 5. Pairwise alignments between PopAlign models of Mammary Gland and Limb Muscle can be dissected in terms of $\Delta w$, $\Delta \mu$, $\Delta \Sigma$** Aligned subpopulations between the reference tissue, Mammary Gland, and the test tissue, Limb Muscle, are ranked by their Jeffrey's Divergence (a). For each aligned pair, we display: (b) associated p-value, (c) mean gene expression states ($\mu_i$) in terms of annotated features, (d) shifts in abundance ($\Delta w$), (e) shifts in mean gene expression state ($\Delta \mu$), (f) shifts in population spread ($\Delta \Sigma$).

**SI Figure 6. Change in abundance for T cells and Monocytes across all drugs** (a) Shifts in abundance ($\Delta w$) for monocyte populations in each drug sample that align to monocytes from the control. (b) Shifts in abundance ($\Delta w$) for T cell populations in each drug sample that align to T cells from the control. Drugs are organized in terms of increasing $\Delta \mu$ so that they share the same ordering as Figure 5c-d.

**SI Figure 7. t-SNEs for all drugs** t-SNE plots for all tested drugs. In each plot, cells for that specific sample are highlighted in purple, while cells from the entire experiment are plotted as gray dots in the background. Glucocorticoids, mTOR inhibitors, and the prostaglandin (alprostadil) are highlighted in magenta, blue, and green.

**SI Table 1. Patient information for PBMC donors** Patient information table with Age, Gender, Ethnicity, Disease Status, and Medications.

**SI File 1. Excel spreadsheet giving cell-type specific differentially expressed genes for all drug samples.** Sheet 1 (degenes_by_celltype): Number of genes and gene names by drug and cell type. Sheet 2 (degenes_percent): percentage of differentially expressed genes.
